# Differential routing of spectral light inputs redistributes circadian phase sensitivity

**DOI:** 10.64898/2026.08.07.743459

**Authors:** Quentin Thommen

## Abstract

Shared sensory channels create a routing problem when biological functions require different combinations of the same environmental inputs. In phototrophic systems, spectral light signals can influence both circadian entrainment and irradiance-dependent physiology. We formulate a two-channel framework that separates common changes in channel activity from differential changes in their relative balance and asks how a locked oscillator represents these directions. Input routing and entrainment stability are inseparable in the locked state: common and differential sensitivities depend on signed, cycle-averaged pathway projections together with the local stiffness of locking. Temporal gating provides one constructive route to a differential projection because two positive pathways can weight opposite-signed regions of a phase-response curve at different times. A canonical repressilator establishes feasibility, while a distinct zeitgeber-gated nonlinear implementation reproduces the same qualitative directional hierarchy. A prescribed positive physiological projection shows that irradiance responsiveness can remain available to a second output. The resulting principle is a redistribution of sensitivity across input directions rather than global attenuation, with experimentally testable predictions for shared multichannel sensing.

## 1. Introduction

Biological sensory systems often distribute the same environmental signals to several downstream functions. Different outputs need not use the same combinations of those signals, so functional separation must arise downstream from sensing. The resulting routing problem is simple to state: one output may need to attenuate an input component that another output must retain. The key issue is therefore not which channels are sensed, but how their activities are projected onto different outputs.

Light sensing in phototrophic organisms provides a concrete example. Multiple photosensory pathways contribute to light perception across cyanobacteria, algae, and plants [18, 4, 14, 17, 22], while circadian timing remains closely integrated with metabolism in photosynthetic lineages [5, 3]. Environmental light varies along at least two useful local directions. Scaling several spectral channels together changes irradiance while approximately preserving their relative balance; redistributing light among channels changes spectral composition more strongly than total intensity. Depth, optical constituents, weather, and vertical mixing generate mixtures of these components in aquatic environments [19, 24, 11, 10]. Mammalian studies likewise show that spectral composition can modify circadian alignment independently of irradiance, particularly around twilight [35, 25]. Common irradiance and spectral contrast there-fore provide biologically motivated coordinates in input space without implying a universal sensory mechanism.

Circadian timing and irradiance-dependent physiology place different demands on these directions. Day-to-day irradiance changes alter available energy but need not provide useful timing information. A locked clock may therefore benefit from weak sensitivity to a common scaling of active channels while retaining sensitivity to another direction in the same input space. The question concerns redistribution of sensitivity, not an assumption that spectral contrast is intrinsically a better zeitgeber. Previous work established robust entrainment under daylight fluctuations and photoperiodic variation [32, 29, 30, 28], while multi-input models showed how parallel light pathways can expand entrainment and photoperiodic responses [33, 7]. A biologically motivated mathematical model of blue- and green-light entrainment in *Ostreococcus tauri* incorporated LOV-HK and Rhod-HK photosensory pathways into the

TOC1–CCA1 clock network [31]. Multiple pathways, however, do not by themselves determine whether their signed, cycle-averaged phase contributions oppose or reinforce one another.

Several architectures can solve the routing problem. Complete pathway segregation assigns different inputs to different outputs; receptor-level opponency or inputs acting on different oscillator variables can also generate signed opposition. The present study asks a narrower question: can two non-negative pathways acting on the same oscillator variable acquire different cycle-averaged phase projections solely because they access the oscillator at different times? A two-channel phase reduction first separates common and differential sensitivities and exposes the role of locking stiffness. A canonical repressilator then tests whether temporal gating can realize the required signed geometry, while gain, gate-geometry, receptor, and zeitgeber-gated controls delimit that construction. Both effective channels share the same normalized light–dark waveform in the numerical test bed, so the spectral interpretation is one application of a more general multichannel geometry rather than a simulation of daily spectral dynamics in a particular organism.

The routing principle extends beyond circadian photoreception. Any dynamical regulatory system that receives several sensory channels while downstream functions require different combinations of their activities faces the same input–output problem. Temporal filtering already provides selective control of cellular responses in signaling cascades [21]; the present mechanism differs by acting through signed, cycle-averaged projections onto the locked phase of an oscillator. Circadian entrainment offers a biologically grounded setting in which this directional sensitivity can be derived and tested constructively, without assuming that the same mechanism has been demonstrated in other oscillator classes.

## 2. Differential routing of shared sensory inputs

### 2.1. Common irradiance and spectral contrast require distinct output projections

Shared receptor signals can feed both circadian and physiological outputs while each branch combines them differently (Fig. 2). Let *I*_1_(*t*) and *I*_2_(*t*) denote the two spectral irradiances, with monotone receptor responses *S*_*i*_(*I*_*i*_). In the circadian branch, a non-negative gate *G*_*i*_(*θ*) sets when channel *i* reaches the oscillator, while the phase-response projection *Z*_*i*_(*θ*) sets the sign of the phase effect. Their product

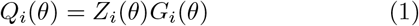

separates pathway activity from signed phase action. Positive gates can therefore generate opposite cycle contributions by sampling opposite signs of the phase-response curve.

**Figure 1:**
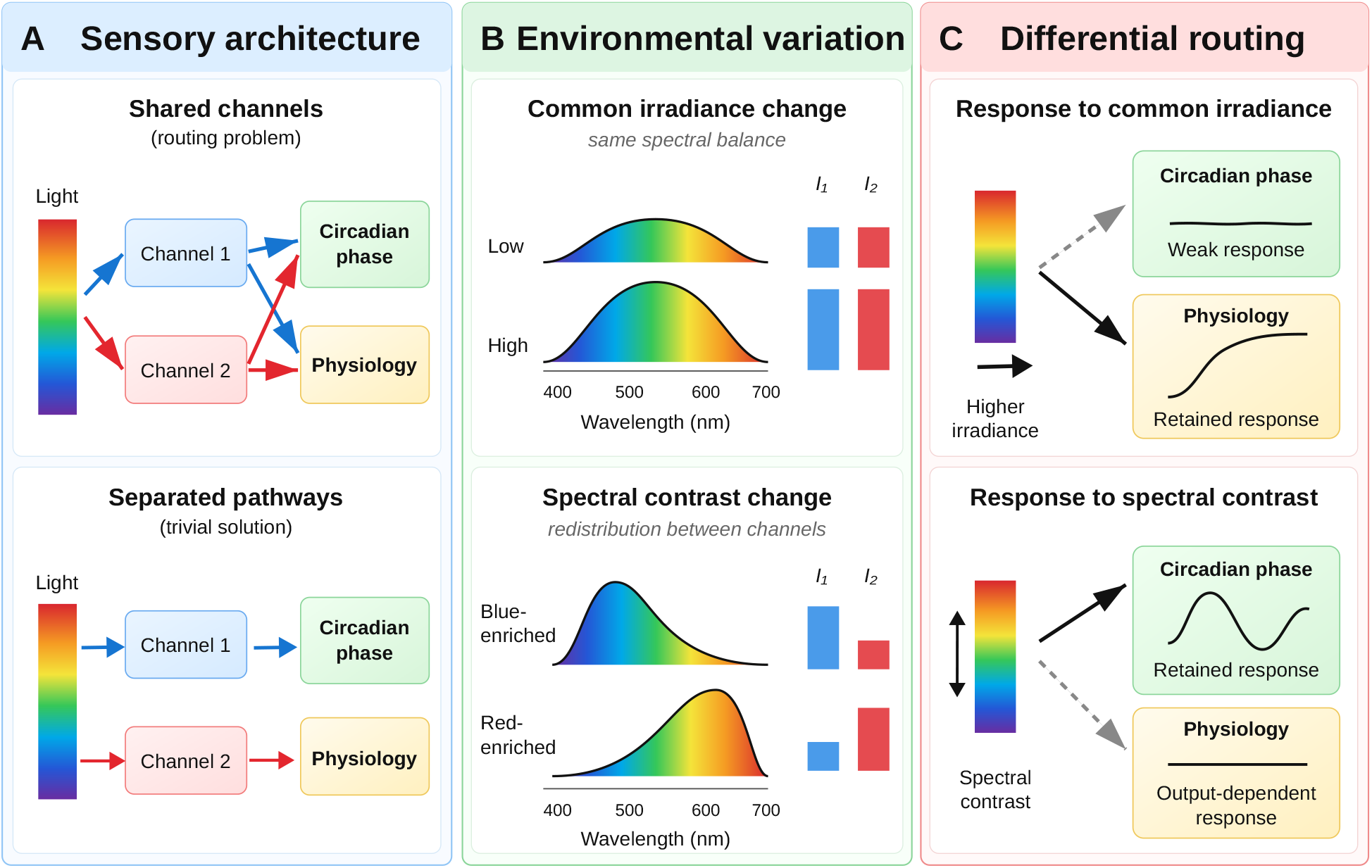
Conceptual overview of differential routing of shared spectral inputs. (A) Multiple sensory channels can contribute to the same downstream functions, creating a routing problem when these functions require different combinations of the inputs. Complete pathway segregation provides an alternative architecture in which distinct channels feed distinct functions. (B) The spectra schematically represent two local directions in input space rather than unique environmental variables: a common irradiance change scales both spectral channels in the same direction while approximately preserving their relative balance, whereas a spectral-contrast change redistributes activity between channels. (C) Differential routing represents an output geometry in which locked circadian phase is less sensitive to the common direction than to the differential direction, while a distinct physiological projection can remain irradiance-responsive. The physiological branch is prescribed as an independent output projection rather than predicted by the oscillator model, and its response to contrast is not constrained by the routing condition.

**Figure 2:**
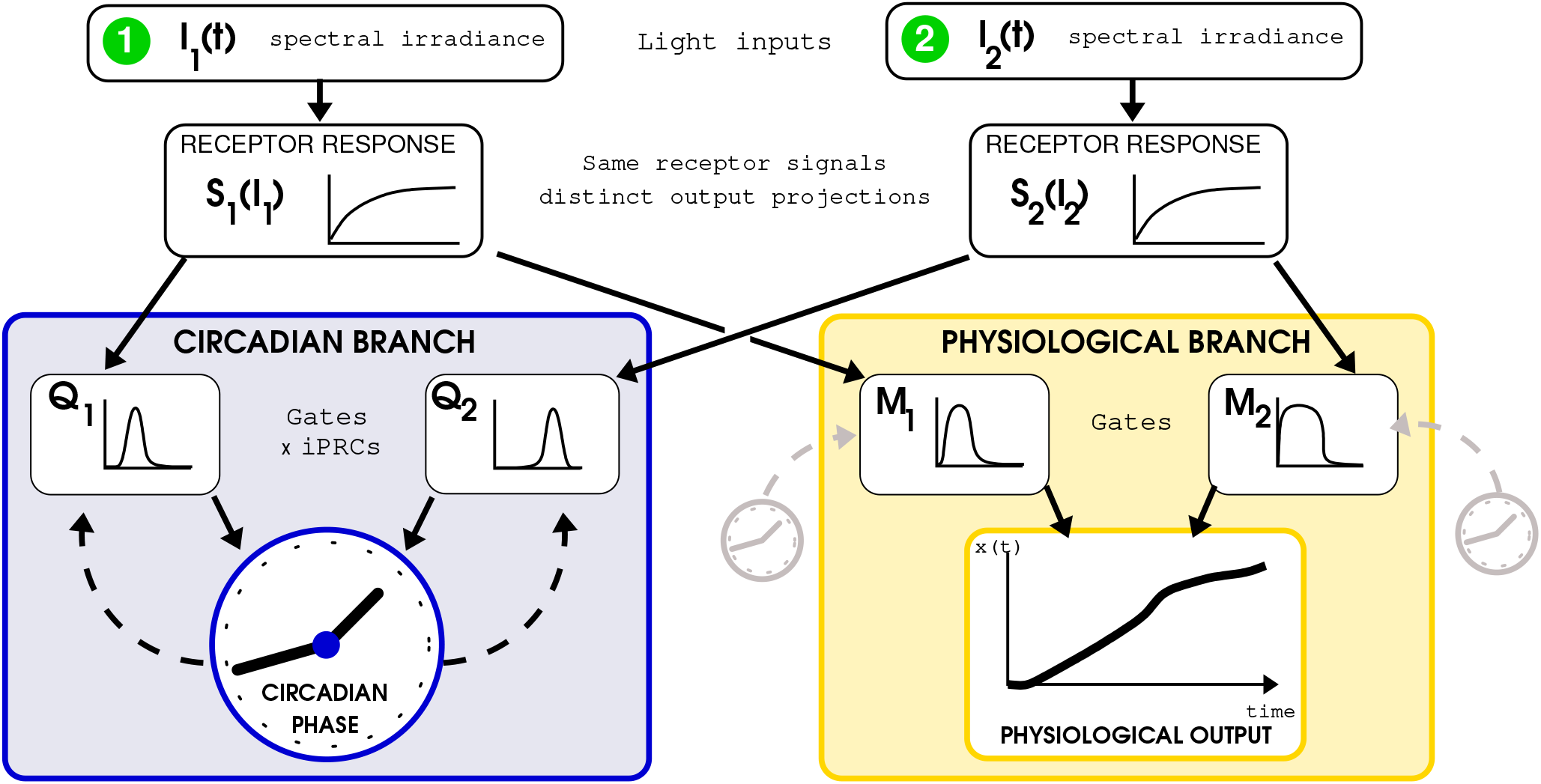
Shared spectral inputs are separated by output-specific projections rather than by pathway segregation. The spectral irradiances *I*_1_(*t*) and *I*_2_(*t*) are transformed by receptor responses *S*_1_(*I*_1_) and *S*_2_(*I*_2_) and remain available to both branches. In the circadian branch, non-negative temporal gates *G*_1_ and *G*_2_ determine when each channel reaches the oscillator; the signed phase effects are *Q*_*i*_ = *Z*_*i*_*G*_*i*_. In the physiological branch, the same receptor outputs are projected through schematic sensitivities *M*_1_ and *M*_2_; these *M*_*i*_ are not simulated in the repressilator model. Differential routing refers to this shared-input architecture.

Channel and gate denote functional roles rather than unique molecular entities. A channel may represent a photoreceptor, an antenna-weighted spectral band, or a broader photosensory pathway; a gate may arise from rhythmic receptor abundance, phase-dependent signal transduction, or downstream access to clock components. The relevant property is the input–output organization: circadian and physiological projections can differ even when they receive the same receptor signals.

Common and differential input directions become explicit in logarithmic coordinates,

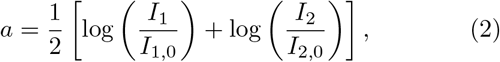

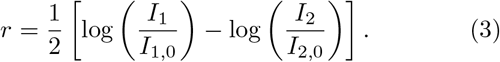

Here *I*_*i*,0_ is the reference irradiance of channel *i, a* scales both channels together, and *r* changes their ratio at fixed geometric mean of the normalized irradiances. Spectral contrast therefore denotes a local direction in two-channel input space rather than a unique physical measure of spectral composition. Appendix A gives the full derivation.

### 2.2. Locked-phase sensitivities decompose into common and differential projections

Locked-phase sensitivity follows directly from the cycle-averaged channel contributions. Let *A*_*i*_ denote the signed contribution of channel *i* at the stable locked phase *ψ*\*. Implicit differentiation of the locking condition gives

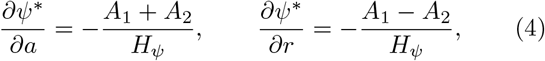

where *H*_*ψ*_ is the local slope of the interaction function. Weak common-mode displacement with retained differential sensitivity therefore corresponds, to first order, to

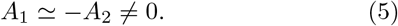

Equation (5) characterizes the local locked-state response; it does not define an independent design rule in every receptor model.

Linear receptors expose the role of entrainment stability. For *S*_*i*_(*I*_*i*_) = *I*_*i*_, a common multiplicative perturbation gives *η*_*i*_ = *∂S*_*i*_*/∂* log *I*_*i*_ = *S*_*i*_, hence

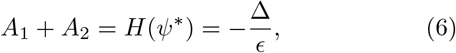

and

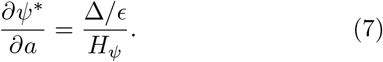

Near resonance, common-mode sensitivity is therefore set by detuning and locking stiffness, while the differential numerator *A*_1_ *A*_2_ remains sensitive to pathway timing and signed phase projection. Saturating receptors remove the exact identity because *η*_*i*_ ≠ *S*_*i*_, but Eq. (4) still couples common-mode sensitivity to both the summed receptor-weighted contributions and *H*_*ψ*_.

A separate physiological projection can use the same receptor signals differently. The present model prescribes this branch: positive contributions of both channels preserve common-mode irradiance responsiveness while the circadian locked state can remain less sensitive to that direction. The analysis therefore addresses compatibility between distinct output projections rather than emergence of the physiological branch.

Global receptor saturation provides a different limit. Vanishing logarithmic receptor gains weaken common-mode, differential, and physiological responses together. Differential routing instead keeps the sensory channels active and changes their signed projection onto locked circadian phase.

### 2.3. Temporal gating can maintain a signed differential contribution from positive pathways

A sign-changing phase-response curve lets the same positive pathway advance the clock at one phase and delay it at another. Two non-negative gates that weight different response windows can therefore generate cycle-averaged contributions of opposite sign. In the realization below, both channels act on the same oscillator variable, so their instantaneous projections share the same sign whenever the gates overlap; the opposition emerges only after temporal weighting and cycle averaging.

Contribution mismatch provides a local description of this geometry. Writing *A*_2_ = −(1 + *δ*)*A*_1_ gives

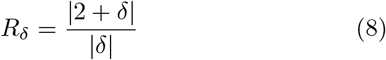

for the ratio of differential to common numerators. A 10% mismatch gives *R*_*δ*_ *≃* 21 and a 20% mismatch gives *R*_*δ*_ *≃*11. These values apply at a fixed locked state. A gain perturbation generally shifts *ψ*\* and changes both *H*_*ψ*_ and the *A*_*i*_, so *δ* cannot be identified with an imposed gain ratio.

Temporal gating thus provides one sufficient route to a large differential component *A*_1_ − *A*_2_. Receptor-level opponency, inputs acting on different oscillator variables, or other architectures can produce the same signed geometry by different means. The nonlinear construction below tests whether temporal separation alone can realize it in a canonical oscillator.

## 3. A nonlinear oscillator provides a constructive test bed

A symmetric three-node repressilator provides a deliberately constructed nonlinear test bed. The model supplies a stable limit cycle, a sign-changing phase-response curve, and controllable temporal access of each effective sensory pathway; it is not a molecular model of a phototrophic clock. Cyclic inhibitory oscillators of this type provide a canonical minimal model of self-sustained biological oscillation [12]. Both effective pathways act positively on the same oscillator variable. Redundant routing gives them the same temporal gate, whereas differential routing assigns temporally separated gates that weight opposite-signed response windows. The second pathway gain follows an unweighted signed-area balance at the reference condition. Both effective channels share the same normalized light profile *L*(*t*) and differ only through the logarithmic amplitudes *a* ± *r*, isolating temporal organization from receptor sign and channel-specific daily waveforms.

### 3.1. A sign-changing phase response provides the substrate for a differential projection

The autonomous repressilator generates a stable limit cycle and an *x*_1_-directed infinitesimal phase-response curve with broad positive and negative lobes (Fig. 3B,C). Temporally separated positive pathways can therefore acquire opposite cycle-averaged signs by weighting different phases of the same iPRC. Because both pathways act positively on *x*_1_, receptor-level opponency plays no role in this sign difference.

**Figure 3:**
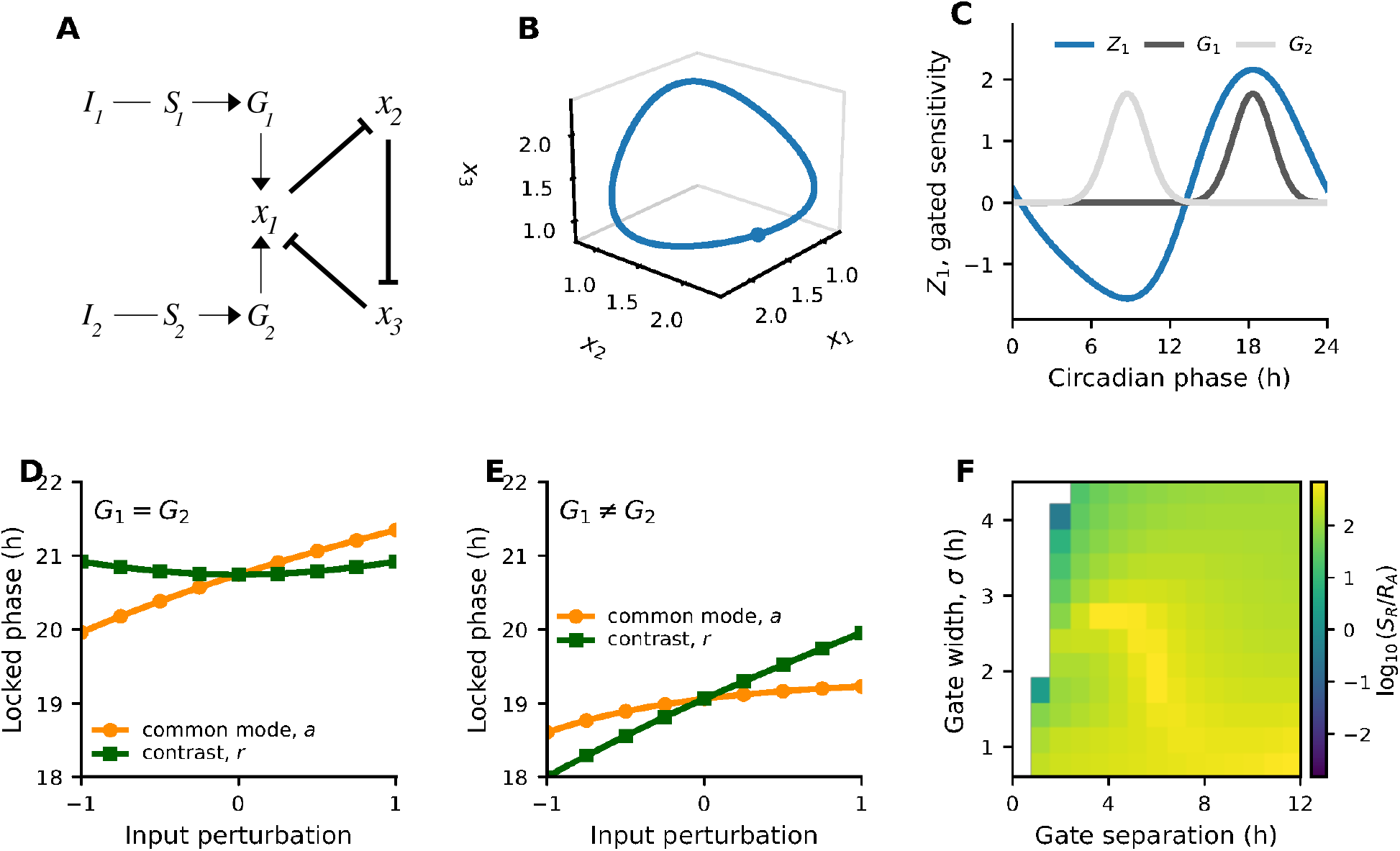
Temporal gating redistributes locked-phase sensitivity between common and differential input directions. (A) Conceptual routing scheme. (B) Stable limit cycle of the autonomous repressilator (*α* = 5, *n* = 3). (C) *x*_1_-directed iPRC *Z*_1_ and two non-negative gates placed on temporally separated response lobes. (D) Redundant routing, *G*_1_ = *G*_2_, is primarily common-mode sensitive in the representative locked state. (E) Differential routing, *G*_1_ ≠ *G*_2_, is more sensitive to the contrast direction. (F) Regularized selectivity Σ = log_10_[(*S*_*R*_ + *ϵ*_0_)*/*(*R*_*A*_ + *ϵ*_0_)] across gate separation and width at zero detuning, with the relative gain rebalanced for each geometry. Blank cells indicate configurations that could not be balanced or lacked a stable reference phase. Panel F shows that the construction is not restricted to one gate pair; it is a feasibility map rather than a robustness map for a fixed architecture. Panels D–F were computed from the averaged interaction function.

Representative gate centres map to approximately ZT23.26 and ZT13.68 in the locked reference state, where the normalized light levels are about 0.17 and 0.026. Effective forcing consequently comes mainly from gate tails that overlap the rising and falling light transitions. The representative construction is therefore dawn/dusk-weighted. That feature belongs to this test bed and makes photoperiod and light-profile shape relevant to quantitative interpretation; the general two-channel framework does not require it.

### 3.2. Differential gating redistributes locked-phase sensitivity toward the contrast direction

Redundant gating produces predominantly commonmode sensitivity. Around the reference condition, the locked phase changes by approximately 0.68 h per logarithmic unit of common irradiance and 0.01 h per logarithmic unit of spectral contrast (Fig. 3D). Across the full perturbation interval, common-mode perturbations shift the locked phase by about 1.39 h, whereas contrast produces little displacement.

Differential gating reverses this hierarchy. Temporally separated gates increase the signed differential component sampled over the cycle, yielding local sensitivities of approximately 0.27 h per logarithmic unit for common irradiance and 0.96 h per logarithmic unit for contrast (Fig. 3E). Across the same perturbation range, the locked phase varies by approximately 0.62 h along the common direction and 1.96 h along the contrast direction. The local contrast response is therefore about 3.6 times larger than the residual common-mode response.

The numerical comparison demonstrates redistribution of locked-phase sensitivity rather than direct pathway cancellation. Equation (4) couples *R*_*A*_ to both *A*_1_ + *A*_2_ and *H*_*ψ*_, while Eq. (7) further constrains the common-mode numerator for linear receptors. The architecture-specific result is therefore the retained differential response and the hierarchy *S*_*R*_ *> R*_*A*_. The representative gates establish feasibility; the scans below test how that construction behaves when gain and gate geometry vary.

### 3.3. A qualitatively similar hierarchy appears in a distinct nonlinear implementation

Direct simulations tested a second implementation of temporally separated access. The averaged model ties gates to oscillator phase, whereas the complete periodically forced repressilator fixes them in zeitgeber time. The direct simulation therefore tests a distinct nonlinear timing rule rather than a nonlinear validation of the phase-gated equations.

Saturating receptors gave local sensitivities of 0.58 h per logarithmic unit to common irradiance and 0.06 h per logarithmic unit to contrast under redundant routing, versus 0.40 and 2.36 h per logarithmic unit under differential routing. Differential selectivity therefore reaches about 5.9 in the direct implementation, compared with about 3.6 in the averaged representative state. Linear receptor responses preserved the same ordering, showing that receptor saturation is not required for the qualitative hierarchy. In the linear case, Eq. (7) also constrains common-mode sensitivity.

Gain variation tests the representative phase-reduced architecture at fixed detuning (Fig. 4). Stable locking persists across all 41 values of *γ/γ*\* = 0.5–1.5. Spectral-contrast sensitivity varies only from 0.932 to 0.965 h per logarithmic unit, while common-mode sensitivity spans 0.202–0.376 h per logarithmic unit; *S*_*R*_*/R*_*A*_ therefore remains between 2.48 and 4.79. The scan shows continuity of the locked solution and persistence of the directional hierarchy over the tested range. It does not test Eq. (8), because *γ/γ*\* is a pathway-gain ratio rather than the contribution mismatch *δ*, and changing gain moves the locked phase.

**Figure 4:**
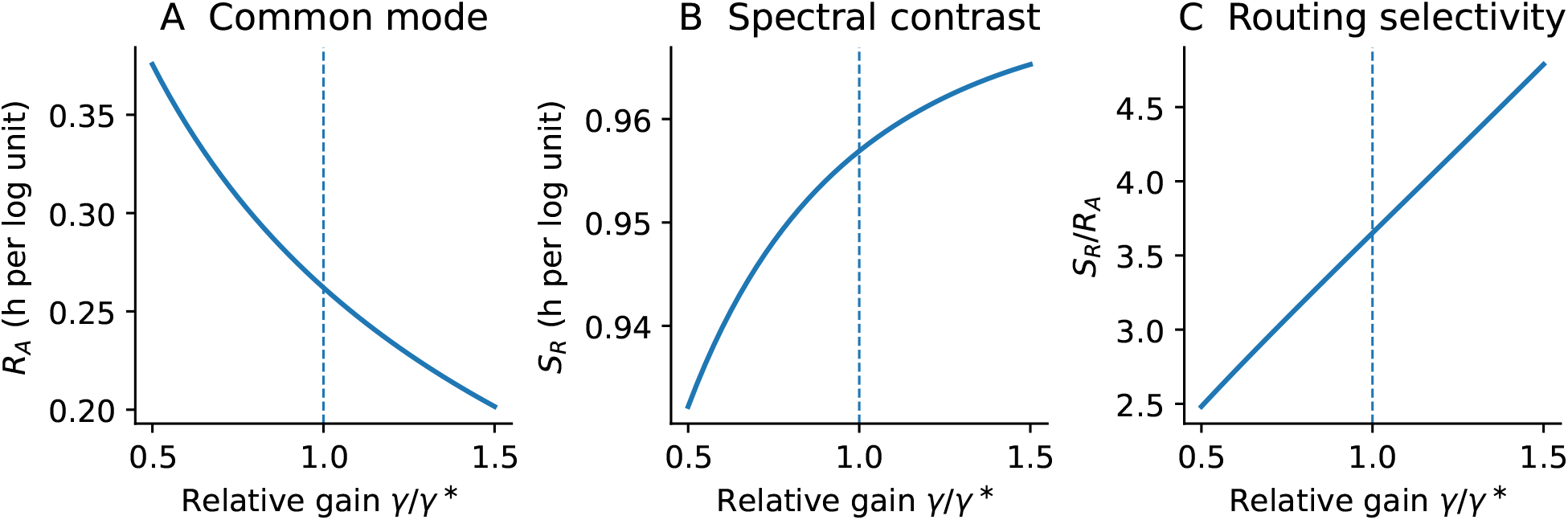
Response of the representative locked state to variation of relative pathway gain. (A) Local common-mode sensitivity *R*_*A*_, (B) spectral-contrast sensitivity *S*_*R*_, and (C) their ratio *S*_*R*_*/R*_*A*_ as functions of *γ/γ*\*. The detuning and gate geometry were kept fixed. Stable locking was retained over *γ/γ*\* = 0.5–1.5, and the hierarchy *S*_*R*_ *> R*_*A*_ persisted throughout the scan. The vertical dashed line marks the unweighted signed-area balance *γ* = *γ*\* used to construct the reference architecture; it is not a numerically optimized gain and *γ/γ*\* should not be identified with the contribution mismatch *δ* of Eq. (8).

Weak variation of *S*_*R*_ across the gain scan is consistent with relocation of the locked state rather than fixed cycle-averaged contributions. The reference value *γ*\* denotes the unweighted signed-area balance used to construct the architecture; it is neither a numerical optimum nor a measured cancellation point.

### 3.4. Contrast-responsive locked states exist across a finite range of temporal geometries

Gate-geometry scans test whether the differential response depends on one specific pair of gate positions (Fig. 3F). A broad set of admissible configurations retains both stable locking and non-zero differential sensitivity.

The feasibility map uses zero detuning and rebalances relative gain at each geometry. For linear receptors, zero detuning forces the common-mode numerator to vanish at a locked state [Eq. (6)], so large selectivity values near can-cellation do not measure robustness or biological optimality. The map supports a narrower conclusion: temporally separated gates can sustain differential sensitivity across a finite range of separations and widths. Small separations make the pathways functionally similar, whereas broad gates average over positive and negative response regions and reduce the differential component.

Geometry and gain scans therefore probe different properties. The geometry map tests existence across rebalanced temporal organizations; Fig. 4 follows one fixed geometry while pathway strength changes. Neither analysis provides a joint robustness map.

## 4. Functional and experimental consequences

### 4.1. Functional compatibility illustration

A minimal light–capacity model illustrates one consequence of the different locked-phase sensitivities without testing adaptive value. The energetic branch responds positively and identically to irradiance in both routing architectures, while rhythmic photosynthetic capacity is prescribed independently of the routing mechanism. Day-to-day irradiance fluctuations therefore alter available energy but carry no additional timing information in this construction. Common-mode variability displaces the clock in both architectures, with systematically smaller excursions under differential routing (Fig. 5B). At the largest variability tested, mean absolute phase displacement reaches approximately 0.254 h with redundant gating and 0.113 h with differential gating. The reference profiles in Fig. 5A show the corresponding overlap of light, photosynthetic capacity, and gross production.

**Figure 5:**
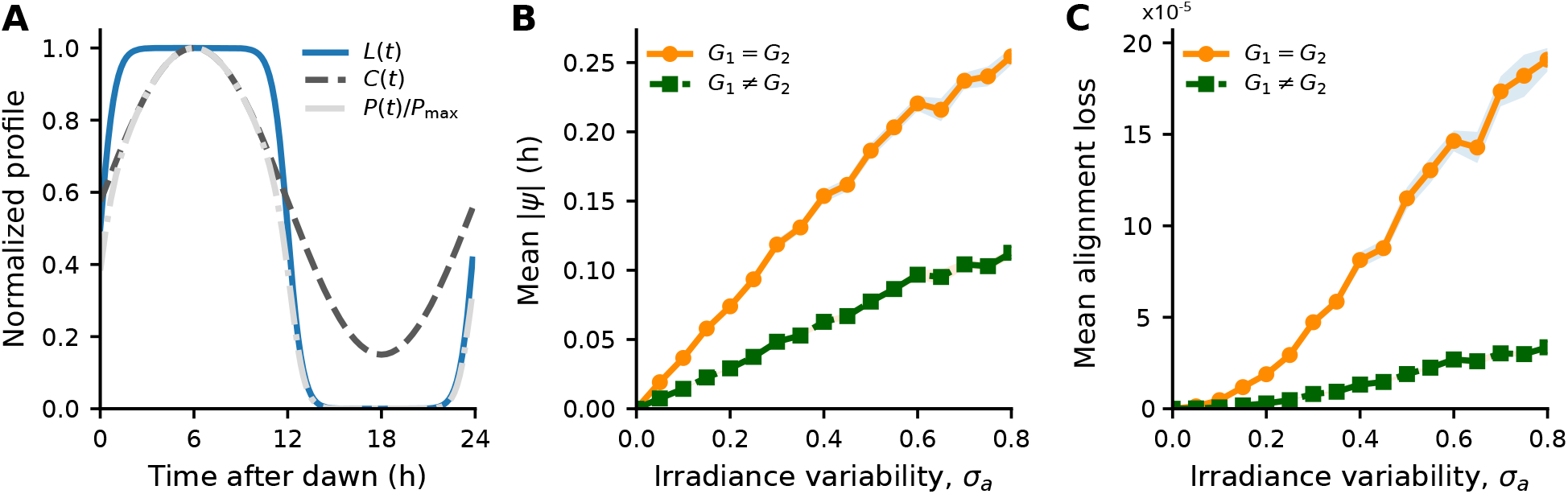
Functional illustration of light–capacity alignment. (A) Normalized reference profiles illustrating the temporal overlap between light availability, photosynthetic capacity, and gross production. (B) Mean absolute phase displacement versus the standard deviation *σ*_*a*_ of day-to-day common-mode log-irradiance fluctuations. (C) Mean alignment loss, *⟨L⟩* = *⟨*Λ^(0)^ *−* Λ*⟩*, where Λ^(0)^ is evaluated for the same daily irradiance realization with the capacity phase held at its reference value. Both architectures retain the same prescribed positive energetic response; differential routing reduces phase displacement and the resulting alignment penalty. At the largest irradiance variability, the alignment-loss quantity is reduced by about 82%; under the same parameterization the difference in mean net production is approximately 0.104%. Lines show means over paired realizations and shaded regions show standard errors.

Reduced phase excursions improve light–capacity alignment. At the largest irradiance variability, mean alignment loss falls from approximately 1.91 × 10^*−*4^ in the redundant architecture to 3.36 × 10^*−*5^ under differential routing, a reduction of about 82% (Fig. 5C).

Mean production changes much less than the alignment-loss metric. At the largest variability, the difference is approximately 1.57 × 10^*−*4^, or about 0.104% of the redundant architecture’s mean net production. The functional result is therefore limited to compatibility: reduced phase displacement can lower an alignment penalty while irradiance remains available to the energetic branch.

### 4.2. Matched common and contrast perturbations test the routing geometry

Matched multichannel perturbations can test the routing geometry without reconstructing the complete molecular network. A common-mode perturbation changes both effective channels by the same logarithmic increment, whereas a contrast perturbation changes them in opposite directions from the same entrained reference state. Experimental channel coordinates should follow receptor action spectra or an appropriate mixing matrix rather than narrow spectral bands by assumption. Conservation of total photon flux does not in general coincide with the model coordinate *r*, especially when baseline channel activities differ.

The ratio *S*_*R*_*/R*_*A*_ provides a direct quantitative read-out. Values above unity indicate greater local locked-phase sensitivity to the differential direction than to the common direction; stronger thresholds should reflect measurement noise and biological effect size. Independent monochromatic response curves cannot recover this joint input geometry.

A second output distinguishes output-specific routing from global sensory attenuation. The present framework assigns physiological irradiance responsiveness through a separate positive projection. Experimentally, a two-dimensional response surface for photosynthetic activity, acclimation, biomass accumulation, or growth can test whether that output retains a common-mode component while locked circadian phase remains less sensitive to it.

Slow or cycle-to-cycle perturbations probe the locked-state geometry developed here. Impulses at a fixed phase address a different observable and can produce large transient shifts even when cycle-averaged locked-state sensitivity is small.

### 4.3. Channel-resolved timing tests the temporal-gating hypothesis

Selective pathway perturbations distinguish the multi-channel requirement from the temporal mechanism. Removing one pathway or changing its gain perturbs any opponent architecture, whereas channel-resolved phase-response measurements or shifts of one temporal access window directly test temporal gating. Reducing temporal separation should reduce the differential projection if the proposed mechanism operates.

Matched common/contrast perturbations and timing perturbations therefore address two levels of inference. The former test output geometry; the latter test the temporalgating hypothesis. A unique molecular architecture requires additional system-specific evidence.

Photoperiod remains a major boundary condition for the representative implementation because effective forcing comes mainly from gate tails overlapping dawn and dusk. Changes in photoperiod or light-transition shape alter those overlaps and therefore modify both *A*_*i*_ and *H*_*ψ*_. Preservation of the same directional hierarchy across photoperiods would require compensating changes in pathway timing, gain, phase sensitivity, or other coupling features. The present study does not test that dependence.

## 5. Discussion

Shared sensory channels can support different output requirements only if downstream dynamics project them differently. We addressed this problem at three levels: a two-channel phase reduction for locked-state sensitivities, a repressilator construction testing temporally separated positive pathways, and a prescribed energetic branch illustrating compatibility with a second light-dependent output. Gain, gate-geometry, receptor, and zeitgeber-gated controls delimit the numerical construction. The combined analysis shows that directional routing in a locked oscillator depends jointly on signed pathway projections and on the local stiffness of entrainment.

The analytical decomposition separates input geometry from the stability of locking. Common and differential sensitivities depend on the summed and differential pathway contributions relative to *H*_*ψ*_. Opposing cycle-averaged contributions therefore characterize weak common-mode displacement with retained differential sensitivity, but they do not constitute an independently tunable design condition in every receptor model. Linear receptors make the constraint explicit: under common multiplicative forcing, the locking equation fixes the summed contribution through detuning, so common-mode sensitivity is controlled by detuning and locking stiffness. Pathway organization remains free to shape the differential contribution.

Temporal gating supplies one constructive route to that differential component. Two positive pathways acquire opposite cycle-averaged signs when they weight different phases of a sign-changing PRC. In the same-state-variable repressilator, the pathways retain the same instantaneous sign wherever their gates overlap; opposition appears only after temporal weighting over the cycle. Receptor opponency, chromatic adaptation, inputs acting on different oscillator variables, or other network mechanisms could generate the same signed geometry by different means [18]. Temporal filtering in signaling cascades already supports selective downstream responses [21]; the specific contribution here is the formulation of routing as a directional redistribution of locked-phase sensitivity in multichannel input space, together with a constructive temporal-gating realization using two positive pathways.

Previous circadian models clarify the scope of that contribution. Robust entrainment and PRC constraints were established in single-input settings [32, 29, 28], while multi-input models showed how parallel light pathways expand entrainment and photoperiodic flexibility [33, 7]. The published *Ostreococcus tauri* model likewise incorporated multiple photosensory contributions in a biologically motivated circadian network [31]. Such architectures establish the relevance of multi-input entrainment but do not establish the signed projection geometry studied here. The present repressilator isolates the narrower mechanism in which temporal separation alone produces opposite cycle-averaged contributions from non-negative inputs acting on the same state variable. Whether *O. tauri* or any other organism implements this geometry remains an experimental question.

The numerical controls support distinct claims. The averaged model demonstrates a reversal from predominantly common-mode to predominantly differential locked-phase sensitivity. The geometry map shows that the construction extends beyond one gate pair, but gain is rebalanced at each point and detuning is zero, so the map addresses feasibility rather than fixed-architecture robustness. The gain scan follows one architecture at fixed detuning and preserves *S*_*R*_ *> R*_*A*_ as the locked state shifts; it does not validate the local mismatch formula. The accompanying SBML models implement a distinct zeitgeber-gated non-linear realization, providing an implementation-level test of the same qualitative directional hierarchy without constituting a strict nonlinear validation of the phase-gated construction.

The representative construction relies strongly on dawn and dusk. Gate centres lie in low-light parts of the cycle, and effective forcing comes mainly from their overlap with the rising and falling light transitions. Photoperiod and transition shape therefore matter quantitatively for this realization. Real spectral composition also changes through the day, and real photoreceptor action spectra overlap, whereas the numerical channels share the same normalized waveform. Spectral contrast should consequently be read as an effective local direction in input space rather than as a literal redistribution between two independent monochromatic sensors.

Differential sensitivity carries a trade-off rather than an automatic biological benefit. A stronger response to contrast is useful only when that direction carries relevant information; environmental noise along the same direction would instead increase vulnerability. Phototrophic systems remain a natural test case because they combine multiple spectral sensors with circadian and metabolic outputs [6, 17, 5, 3, 1, 23], but the specific temporal architecture constructed here remains hypothetical.

The physiological branch serves a narrower purpose. Spectral regulation of photosynthetic acclimation occurs in marine phototrophs, including blue–green light-dependent state transitions in *Ostreococcus tauri* [20] and phytochrome-dependent acclimation in diatoms [9]. These observations motivate a spectrally responsive second output, while the positive projection used here remains prescribed. The light–capacity calculation therefore illustrates compatibility rather than mechanism: reduced phase displacement lowers the alignment penalty under the chosen parameters, whereas mean production changes by only about 0.1%. No generic adaptive or fitness benefit follows from that result.

The framework applies to a stable entrained state under weak, near-resonant forcing and to slowly varying or cycle-to-cycle changes in input amplitude. Fast fluctuations, impulse responses, channel-specific daily spectra, overlapping receptor action spectra, photoperiod dependence, and coupling through different oscillator variables lie outside the present tests. Those limits still leave a direct experimental core: matched common and differential perturbations measure the output geometry, while channel-resolved timing perturbations test whether temporal gating contributes to the observed differential projection.

## 6. Conclusion

Shared sensory channels generate common and differential input directions whose effects on a locked oscillator depend on distinct signed projections. Common-mode displacement follows the summed pathway contribution relative to locking stiffness, whereas differential sensitivity follows the difference between pathway contributions. For linear receptors, detuning also constrains the common-mode component through the locking condition; temporal gating therefore shapes routing without acting as the sole source of common-mode attenuation.

Temporal gating contributes constructively by letting two positive pathways weight opposite regions of a sign-changing phase-response curve and maintain a large differential projection without receptor-level inhibition. A repressilator demonstrates this mechanism in one canonical oscillator, and a distinct zeitgeber-gated nonlinear implementation reproduces the same qualitative hierarchy. Stable locking, signed phase response, and distinct temporal access can occur beyond this topology, although the present study does not test other oscillator classes. A separately prescribed physiological projection shows that irradiance responsiveness can remain available to another output.

The central prediction is experimentally accessible: multiple sensory pathways produce differential routing only when their signed, cycle-averaged projections generate the appropriate input–output geometry. Matched multichannel perturbations probe that geometry, while channel-resolved timing perturbations test the temporal-gating mechanism.

## Data and model availability

### Data and model availability

The SBML implementations of the redundant and differential routing architectures, together with all analysis scripts, parameter files, numerical outputs, and plotting routines required to reproduce the gate-geometry scan, gain scan, direct nonlinear controls, and stochastic light–capacity calculations in Figs. 3–5, accompany the manuscript as Supplementary Material. The SBML models will also be deposited in BioModels upon acceptance.

### Author contributions

Quentin Thommen: Conceptualization, Methodology, Formal analysis, Software, Investigation, Visualization, Writing – original draft, Writing – review and editing.

### Funding

The author received no financial support for the research, authorship, and/or publication of this article.

### Declaration of generative AI and AI-assisted technologies in the manuscript preparation process

During preparation of this work, the author used OpenAI’s ChatGPT to assist with manuscript organization, language editing, code drafting, and numerical workflow development. All generated material, code, parameter choices, numerical procedures, and outputs were reviewed and validated by the author, who takes full responsibility for the content of the publication.

## Appendix A. Analytical derivation of the routing conditions

This appendix gives the phase-reduction and averaging derivation underlying the routing conditions summarized in the main text. The main text retains only the quantities required to interpret the mechanism and the numerical results.

### Appendix A.1. Two spectral inputs and two functional projections

Let *X*(*t*) *∈* ℝ^*n*^ denote an oscillatory regulatory system receiving two spectral irradiances *I*_1_(*t*) and *I*_2_(*t*). Each channel is transformed by a smooth, increasing receptor response *u*_*i*_ = *S*_*i*_(*I*_*i*_), *i* = 1, 2, and the full dynamics is

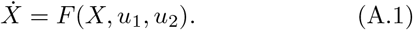

Assume that the reference system has a stable limit cycle *X*_0_(*θ*) with angular frequency *ω*, and let *Z*(*θ*) be its infinitesimal phase-response vector. The signed phase projection associated with channel *i* is

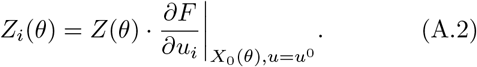

Here, the input derivative is evaluated along the reference limit cycle and at the reference input state *u* = *u*^0^. Temporal access of each channel to the clock is represented by a non-negative gate *G*_*i*_(*θ*) 0. The effective signed phase contribution is *Q*_*i*_(*θ*) = *Z*_*i*_(*θ*)*G*_*i*_(*θ*). This notation separates when a pathway is active from the signed effect of that activity on phase. To first order,

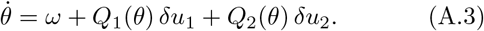

The same spectral inputs can be routed independently toward an energetic or growth-related output. With local sensitivities *M*_*i*_(*θ*), a minimal representation is

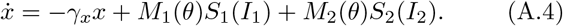

The essential requirement is that (*M*_1_, *M*_2_) need not be proportional to (*Q*_1_, *Q*_2_).

### Appendix A.2. Common-mode and spectral-contrast coordinates

The logarithmic coordinates *a* and *r* are defined by Eq. (3), so that *I*_1_ = *I*_1,0_ exp(*a*+*r*) and *I*_2_ = *I*_2,0_ exp(*a*− *r*). In the symmetric numerical realization, *I*_1,0_ = *I*_2,0_ = *I*_0_.

The logarithmic receptor gains are

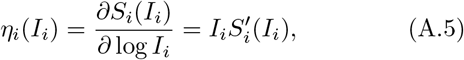

with *η*_*i*_ *>* 0 for active monotone receptor responses. The instantaneous phase sensitivities are

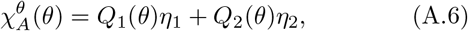

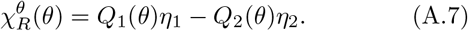

Exact instantaneous rejection of the common mode with retained contrast sensitivity requires

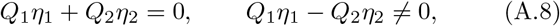

and therefore *Q*_1_*Q*_2_ *<* 0 at that same phase when both channels remain active. This pointwise condition is stronger than the cycle-averaged condition used below. In the same-state-variable repressilator realization, *Q*_*i*_ = *Z*_1_*G*_*i*_ with *G*_*i*_ ≥ 0, so *Q*_1_*Q*_2_ ≥ 0 whenever both gates overlap at the same phase; opposition arises only after the two gates weight different phases over the cycle.

The energetic sensitivity to common irradiance is

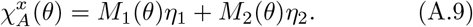

For an energetic output that increases with both spectral inputs,

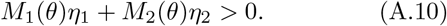

If (*M*_1_, *M*_2_) were proportional to (*Q*_1_, *Q*_2_), common-mode phase cancellation would also cancel the energetic response. In this local representation, retained energetic responsiveness under phase cancellation therefore requires

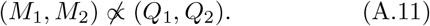

### Appendix A.3. Locked-phase sensitivity

For *T*-periodic forcing, let *ψ* be the phase difference between the oscillator and the external cycle and let Δ denote the frequency mismatch in the rescaled phase equation. With an explicit weak-coupling amplitude *ϵ*,

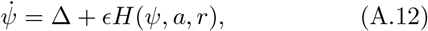

and a stable locked phase *ψ*\* satisfies

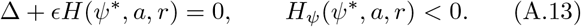

To first order,

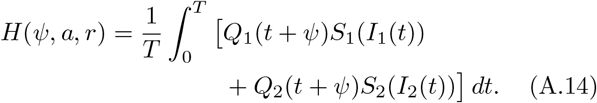

Define the signed cycle-averaged receptor-weighted contributions

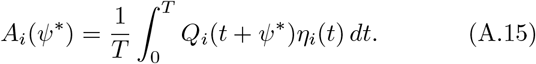

Implicit differentiation gives

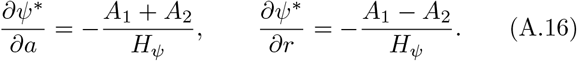

Thus Eq. (5) is, to first order, a characterization of a locked state with weak common-mode sensitivity and retained differential sensitivity. It should not be interpreted as an independent design constraint in every receptor model.

For linear receptors, *S*_*i*_(*I*_*i*_) = *I*_*i*_ and hence *η*_*i*_ = *S*_*i*_.

Under a common multiplicative perturbation,

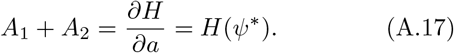

Using the locking condition gives

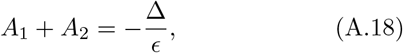

and therefore

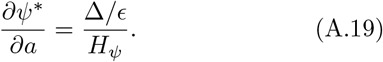

At zero detuning the first-order common-mode sensitivity consequently vanishes for any stably locked architecture satisfying these linear-receptor assumptions, whereas the differential sensitivity remains controlled by *A*_1_ − *A*_2_ and *H*_*ψ*_. For nonlinear receptors such as *S*(*I*) = *I/*(1 + *I*), *η* = *S*(1 − *S*) = *S* and this exact identity no longer holds.

Global receptor saturation provides a different limiting mechanism. If both receptor gains vanish, *η*_1_ *≃ η*_2_ *≃* 0, common-mode and differential phase sensitivities both decrease, together with any physiological response that depends on the same receptor gains.

### Appendix A.4. Local parametrization of contribution mis-match

Let

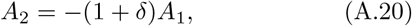

where *δ* is the relative mismatch between the signed cycle-averaged pathway contributions. Then

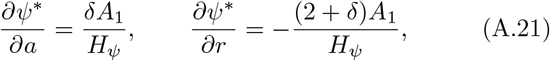

and

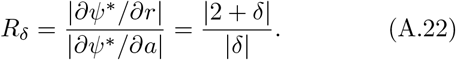

For |*δ*| *≪* 1, *R*_*δ*_ *≃* 2*/*| *δ*|. Thus a 10% mismatch of the cycle-averaged contributions gives approximately 21-fold selectivity and a 20% mismatch approximately 11-fold selectivity at a fixed locked state. This algebraic parametrization does not predict the effect of varying a pathway gain, because gain changes can move *ψ*\* and change both *H*_*ψ*_ and the *A*_*i*_. More generally,

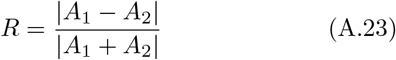

when both sensitivities are evaluated at the same locked state.

### Appendix A.5. Experimental observables

For a short perturbation of channel *i* with duration Δ*t* applied near circadian phase *ϕ*,

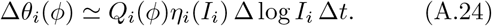

Defining *q*_*i*_(*ϕ*) = *Q*_*i*_(*ϕ*)*η*_*i*_(*I*_*i*_) gives the phase-resolved impulse responses

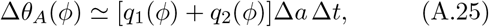

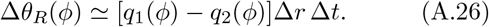

These quantities probe the two pathways at a specified phase and need not cancel pointwise in the temporal-gating architecture studied here. The routing criterion for the *locked phase* instead concerns the cycle-averaged coefficients *A*_*i*_ of Appendix A.3: weak common-mode displacement with retained differential sensitivity corresponds locally to a small *A*_1_ +*A*_2_ relative to *A*_1_ − *A*_2_ and the locking stiffness. Pointwise opposition,

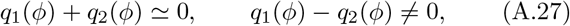

would define a stronger, distinct architecture and is not required by the positive-pathway temporal-gating mechanism simulated here. Phase-resolved measurements of the individual *q*_*i*_(*ϕ*) are nevertheless useful for testing whether the two channels access different signed regions of the response curve.

For acclimated population growth, a two-dimensional response surface *λ*(*I*_1_, *I*_2_) has logarithmic gradient

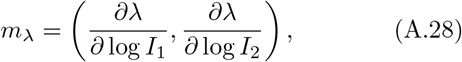

with common-mode and contrast components

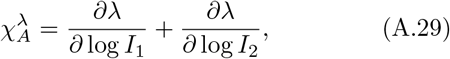

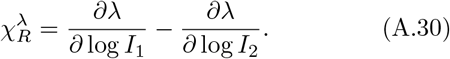

Differential routing requires 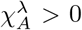 under conditions in which the corresponding circadian common-mode response remains small.

## Appendix B. Numerical implementation and parameter values

OpenAI’s ChatGPT was used to assist code drafting and numerical workflow development. All code, parameter choices, numerical procedures, and outputs used in the analyses were reviewed and validated by the author.

### Appendix B.1. Canonical oscillator and spectral forcing

The autonomous system is the symmetric nondimensional three-node repressilator [12],

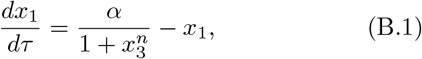

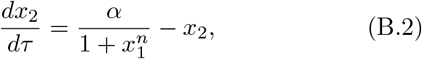

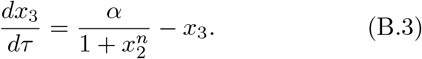

For the weakly forced construction, the input enters additively through the first equation,

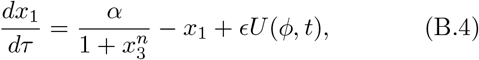

while the *x*_2_ and *x*_3_ equations are unchanged. The oscillator was driven by a smooth 24-h light–dark cycle *L*(*t*). The two effective sensory inputs were parameterized as

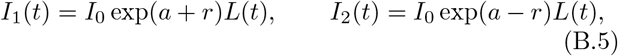

so both channels share the same normalized daily waveform *L*(*t*) and *a* and *r* change only their amplitudes in this test bed. The inputs were transformed by identical saturating receptor responses,

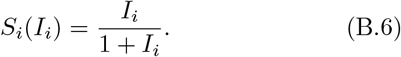

Both pathways acted positively and additively on *x*_1_. Their access to the oscillator was modulated by non-negative periodic Gaussian gates,

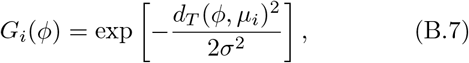

where *d*_*T*_ is the shortest distance on the phase circle. The forcing term was

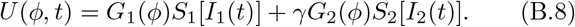

In the redundant architecture, *G*_1_ = *G*_2_. In the differential architecture, the gates were centred on opposite-signed extrema of the *x*_1_-directed iPRC. Their relative gain was chosen from

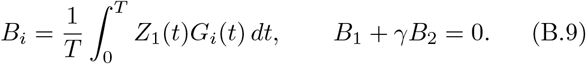

This criterion balances unweighted signed gate areas and therefore does not enforce exact cancellation after daylight weighting and receptor nonlinearity are included.

### Appendix B.2. Phase reduction and selectivity measures

The iPRC was obtained from the adjoint equation with normalization *Z* · *F* = 1 [34, 13, 16, 26]. Under weak coupling,

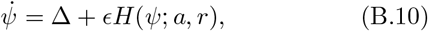

with

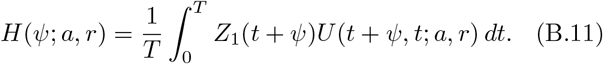

Stable locked phases satisfy

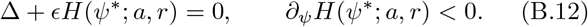

Residual common-mode sensitivity and retained contrast sensitivity were quantified by

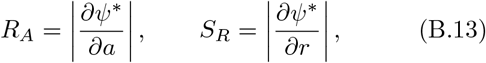

and summarized by

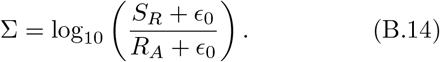

### Appendix B.3. Functional light–capacity model

The energetic contribution of the two spectral channels was represented by

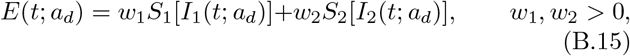

with *r* = 0 in this analysis. Rhythmic photosynthetic capacity was prescribed as

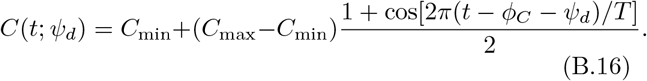

The daily production proxy was

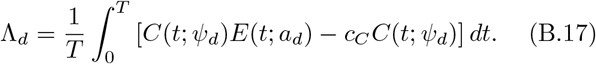

Day-to-day irradiance variability followed

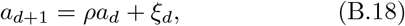

with stationary standard deviation *σ*_*a*_, and phase relaxed toward the irradiance-dependent locked phase according to

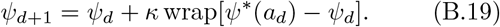

For each day, the reference production 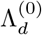 was evaluated under the same irradiance realization *a*_*d*_ and energetic forcing *E*(*t*; *a*_*d*_), with the capacity phase held at its reference value *ψ* = 0. Alignment loss was then defined as

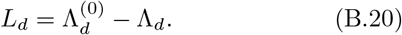

Thus *L*_*d*_ *>* 0 denotes lower production caused by the phase displacement relative to the same-day reference; no rectification is implied by the definition, so a negative value would indicate improved overlap on that day. For each realization, *L*_*d*_ was averaged over the post-burn-in days, and the values plotted in Fig. 5 are means across the paired realizations.

### Appendix B.4. Parameter values and numerical procedures

The autonomous repressilator used *α* = 5 and *n* = 3, giving an autonomous period *T*_0_ = 3.8629206 model-time units. Model time was converted to circadian hours using *t*_*h*_ = 24*τ/T*_0_. The light cycle had period *T* = 24 h and

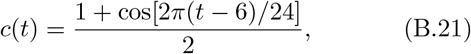

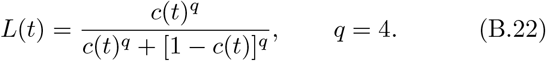

The reference irradiance was *I*_0_ = 1. The representative gates had width *σ* = 1.5 h. In the differential architecture, their intrinsic-phase centres were *µ*_1_ = 18.32 h and *µ*_2_ = 8.74 h, and the relative gain was *γ* = 1.3731. At the reference condition, these corresponded to approximately ZT23.26 and ZT13.68 after locking. With the light profile below, *L*(ZT23.26) *≃* 0.174 and *L*(ZT13.68) *≃* 0.026, so effective forcing is concentrated in the gate tails overlapping the rising and falling light transitions. The redundant gate was centred at approximately ZT21.57.

For the locked-phase curves, *ϵ* = 0.1 and the rescaled phase-equation mismatch was Δ = −0.00290953; with a 24-h forcing period this corresponds to a free-running period of 24.0700 h. The interaction integral used 1200 equally spaced points per cycle; stable roots were detected from a 720-point scan and refined by bracketed root finding.

The gate-geometry map used 15 gate separations between 0 and 12 h and 12 gate widths between 0.6 and 4.5 h. The map was calculated at zero detuning and the relative gain was rebalanced for each geometry. Finite differences used step *h* = 0.08 in the logarithmic coordinates, and the regularization floor was *ϵ*_0_ = 10^*−*3^ h per logarithmic unit. When *R*_*A*_ approaches this floor, Σ is dominated by the regularization and should not be interpreted as the exact logarithm of the unregularized sensitivity ratio.

A gain-imbalance scan was performed on the representative differential architecture by varying the relative gain over *γ/γ*\* *∈* [0.5, 1.5] using 41 equally spaced values, where *γ*\* = 1.37307 is the analytical balance condition of the reference configuration. The detuning Δ was kept fixed at the value used for the locked-phase curves so that changes in oscillator–forcing mismatch did not compensate the imposed gain imbalance. For each gain value, the stable locked phase was computed near *a* = *r* = 0, and the local sensitivities *R*_*A*_ and *S*_*R*_ were estimated with the same centred-difference procedure used for the reference architecture. Stable locking was retained throughout the scan.

The light–capacity model used *w*_1_ = *w*_2_ = 0.5, *C*_min_ = 0.15, *C*_max_ = 1, *c*_*C*_ = 0.12, *ϕ*_*C*_ = 6 h, *ρ* = 0.6, and *κ* = 0.35.

Seventeen values of *σ*_*a*_ were examined between 0 and 0.8. For each value, 25 paired simulations of 160 days were run, with the first 40 days discarded as burn-in.

Direct nonlinear simulations were performed in COPASI using SBML implementations of the forced repressilator. The gates were fixed in zeitgeber time at the reference positions described above. Both saturating *S*(*I*) = *I/*(1 + *I*) and linear *S*(*I*) = *I* receptor responses were tested. Common-mode and contrast perturbations were evaluated over *a, r ∈* [−1, 1]. Simulations continued for 100 days, and local sensitivities were estimated from centred differences around the reference condition.

## Appendix C. Notation

| Symbol | Definition |
|---|---|
| *I<sub>i</sub>*(*t*) | Spectral irradiance in channel *i* |
| *S<sub>i</sub>*(*I<sub>i</sub>*) | Monotone receptor response |
| *X*(*t*) | State vector of the oscillator |
| *Z*(*θ*) | Infinitesimal phase-response vector |
| *G<sub>i</sub>*(*θ*) | Non-negative temporal gate |
| *Q<sub>i</sub>*(*θ*) | Effective signed phase contribution, *Q<sub>i</sub>* = *Z<sub>i</sub>G<sub>i</sub>* |
| *M<sub>i</sub>*(*θ*) | Projection onto a physiological output |
| *a* | Common-mode logarithmic irradiance coordinate |
| *r* | Logarithmic spectral-contrast coordinate |
| *η<sub>i</sub>* | Logarithmic receptor gain |
| *ψ*<sup>\*</sup> | Stable locked phase |
| *A<sub>i</sub>* | Cycle-averaged signed phase contribution |
| *δ* | Relative mismatch between pathway contributions |
| *R<sub>A</sub>* | Common-mode locked-phase sensitivity |
| *S<sub>R</sub>* | Spectral-contrast locked-phase sensitivity |
| Σ | Regularized logarithmic selectivity index |

